# A high-quality *de novo* genome assembly based on nanopore sequencing of a wild-caught coconut rhinoceros beetle (*Oryctes rhinoceros*)

**DOI:** 10.1101/2021.09.12.459717

**Authors:** Igor Filipović, Gordana Rašić, James Hereward, Maria Gharuka, Gregor J. Devine, Michael J. Furlong, Kayvan Etebari

## Abstract

**Background:** An optimal starting point for relating genome function to organismal biology is a high-quality nuclear genome assembly, and long-read sequencing is revolutionizing the production of this genomic resource in insects. Despite this, nuclear genome assemblies have been under-represented for agricultural insect pests, particularly from the order Coleoptera. Here we present a *de novo* genome assembly and structural annotation for the coconut rhinoceros beetle, *Oryctes rhinoceros* (Coleoptera: Scarabaeidae), based on Oxford Nanopore Technologies (ONT) long-read data generated from a wild-caught female, as well as the assembly process that also led to the recovery of the complete circular genome assemblies of the beetle’s mitochondrial genome and that of the biocontrol agent, Oryctes rhinoceros nudivirus (OrNV). As an invasive pest of palm trees, *O. rhinoceros* is undergoing an expansion in its range across the Pacific Islands, requiring new approaches to management that may include strategies facilitated by genome assembly and annotation.

**Results:** High-quality DNA isolated from an adult female was used to create four ONT libraries that were sequenced using four MinION flow cells, producing a total of 27.2 Gb of high-quality long-read sequences. We employed an iterative assembly process and polishing with one lane of high-accuracy Illumina reads, obtaining a final size of the assembly of 377.36 Mb that had high contiguity (fragment N50 length = 12 Mb) and accuracy, as evidenced by the exceptionally high completeness of the benchmarked set of conserved single-copy orthologous genes (BUSCO completeness = 99.11%). These quality metrics place our assembly as the most complete of the published Coleopteran genomes. The structural annotation of the nuclear genome assembly contained a highly-accurate set of 16,371 protein-coding genes showing BUSCO completeness of 92.09%, as well as the expected number of non-coding RNAs and the number and structure of paralogous genes in a gene family like Sigma GST.

**Conclusions:** The genomic resources produced in this study form a foundation for further functional genetic research and management programs that may inform the control and surveillance of *O. rhinoceros* populations, and we demonstrate the efficacy of *de novo* genome assembly using long-read ONT data from a single field-caught insect.

## BACKGROUND

Adult coconut rhinoceros beetles, *Oryctes rhinoceros* L. (Coleoptera: Scarabaeidae), feed by boring into the crown of coconut palms. This damages growing tissue and significantly reduces coconut yields and can lead to the death of trees. Native to southeast Asia, this pest was accidentally introduced into Samoa in 1909 [1], and it has since spread across the tropical Pacific, bringing a significant threat to the livelihoods of the peoples of Pacific island nations for whom the coconut palm (‘the tree of life’) is an important source of food, fibre and timber. Invasive populations of *O. rhinoceros* have been suppressed over the past 60 years through management approaches that included the release of a biocontrol agent, Oryctes rhinoceros nudivirus (OrNV) [2]. However, a highly damaging infestation by *O. rhinoceros* in Guam in 2007 was not controlled with OrNV, and the beetle’s subsequent expansion to other Pacific Islands including Papua New Guinea, Hawaii, Solomon Islands, and most recently Vanuatu and New Caledonia [3]–[6], suggests potential changes in this biological system [7] that require new approaches to management, including the isolation and deployment of highly virulent OrNV strains for specific *O. rhinoceros* genotypes [8].

Genome sequencing has enabled better understanding of population outbreaks, invasion and adaptation mechanisms in insect pests [9], [10]. Functional and comparative genomics studies are identifying new targets for control and the implementation of integrated pest management strategies [11]. Draft genome assembly is generally a good starting point for relating genome function to organismal biology, but the production of this genomic resource for agricultural pests has lagged behind that of some other insects [11], [12]. A recent project aiming to tackle this lag is the Ag100Pest Initiative, led by the United States Department of Agriculture’s Agricultural Research Service (USDA-ARS), that is set to produce reference quality genome assemblies for the top 100 arthropod agricultural pests in the USA, with nearly one third of species belonging to Coleoptera [13].

Draft genome assemblies are very useful for population genomic analyses, enabling the design of, for example, optimal protocols for reduced genome representation sequencing [14]. However, draft genome assemblies that are highly fragmented, incomplete or misassembled have limited use for functional genomic studies. Transcriptome assemblies are useful for studying functionally and sufficiently transcribed parts of the genome, but only complete and accurate genome assemblies provide information on non-transcribed regions (e.g. promoters, enhancers) that can have important influences on gene expression and, ultimately, economically-important phenotypes [13]. In addition, different types of non-translated RNAs (e.g. microRNAs, lncRNAs) are often not detected in transcriptome studies but are included in complete and accurate genome assemblies. These can help us understand how insect pests interact and respond to their hosts, pathogens, the environment and they can reveal new targets for novel genetic control measures (e.g. RNAi [15], gene drives [16], [17]).

Obtaining high-quality genome assemblies is often challenging in insects [12], particularly from short-read sequencing data (e.g. Illumina) for species with high levels of DNA polymorphism and repetitive genomic elements [18]. These issues are further compounded for insects of small physical size or for partial specimens, as they may require whole genome amplification or the pooling of several individuals to obtain enough DNA for library preparations. Different methods of whole genome amplification vary in their ability to preserve specific genetic variation and can be biased against regions with high GC-content, smaller and low-abundance DNA fragments [19]. They can also create chimeric fragments and amplify contaminating DNA that can be erroneously integrated into the target assembly. Pooling individuals is preferably done with individuals from a line that has undergone inbreeding to reduce genetic variation, but many pest species cannot be colonised in the laboratory. Moreover, for those insects that can be lab-reared, intensive inbreeding procedures such as full-sib mating for tens of generations may not reduce heterozygosity in all parts of the genome (e.g. [20]). The pooling of wild-caught samples is particularly problematic given the possibility of combining cryptic species or biotypes, which would impact assembly quality and lead to spurious biological conclusions. When presented with a highly heterozygous genome or a pool of diverse haplotypes, the standard assembly process tends to report a heterozygous region as alternative contigs (instead of collapsing them into a single haplo-contig) and is unable to resolve multiple paths between homo- and heterozygotic regions, producing a highly fragmented assembly with an erroneously inflated total size [21]. Such assemblies cause problems in genome annotation and downstream analyses, giving fragmented gene models, wrong gene copy numbers, and broken synteny. They also preclude linkage mapping and genome-wide association studies.

The development of long-read sequencing technologies is revolutionizing the production of contiguous and complete insect genome assemblies [18], but their requirement for large quantities of input DNA have complicated their application to single-insect assemblies. However, new low-input protocols were recently demonstrated for Pacific Biosciences (PacBio) long-read sequencing, producing high-quality single-insect genome assemblies for the mosquito *Anopheles coluzzii* [22] and spotted lanternfly *Lycorma delicatula* [23]. A chromosome-level assembly was recently reported for a single outbred *Drosophila melanogaster* generated using a combination of long-read sequences from Oxford Nanopore Technologies (ONT), Illumina short-read sequences and Hi-C data [24]. However, the small size of this insect necessitated genome amplification to prepare the sequencing libraries, and the final assembly was ~20% smaller than the canonical reference genome for *D. melanogaster* [24].

Here, we present a high-quality *de novo* genome assembly based on ONT long-read data from a single wild-caught adult female of the coconut rhinoceros beetle (*O. rhinoceros*, NCBI:txid72550). The amount of DNA extracted from this large insect was sufficient to prepare multiple ONT libraries without genome amplification. Data from just one flow cell were enough to produce a high-quality draft assembly of the beetle’s nuclear genome, and data from four MinION flow cells enabled the assembly that is among the most accurate and complete of the published Coleopteran genomes, as well as the assembly of its mitochondrial genome [25], and the genome of the biocontrol agent Oryctes rhinoceros nudivirus (OrNV) [26] that had infected the individual we analysed.

## RESULTS AND DISCUSSION

### Library preparation and sequencing

We used a customized Solid-phase Reversible Immobilization (SPRI) bead-based protocol (see Materials and Methods) to extract high molecular weight (HMW) DNA (Figure 1A-C, Supplemental Figure 1) from an *O. rhinoceros* female caught with a pheromone lure trap in Guadalcanal, Solomon Islands in January 2019. Given the large size of the insect, we achieved high quantity (~10 μg) and quality HMW DNA (Supplemental figure 1) (Figure 1B). We further depleted the low molecular weight (LMW) DNA fragments (<10 kb in length) using the Circulomics XS size-selection kit (Figure 1C), and prepared four standard ligation-based ONT libraries (SQK-LSK109) using a total of 4μg of DNA. Each library was sequenced on a MinION Flow Cell (model R9.4.1, Oxford Nanopore Technologies) (Figure 1D), yielding between 896,000 and 1.48 million raw reads. Using the basecaller Guppy v.3.2.4, we obtained a total 29.4 Gb of sequence data with 89.8% passing the QC filtering (Phred score 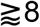). 26.4 Gb of high-quality data with the read length N50 of ~11.3 kb were used for downstream analyses (Supplemental table 1). The longest recorded read was 579.5 kb, but the longest read that passed the QC filtering was 143.6 kb. For the second round of analyses, we used the newer base-caller version, Guppy v4.2.2, that produced a total of 29.5 Gb of data, with 92.1% passing the QC filtering (Phred score 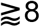). These 27.2 Gb of high-quality reads had a length N50 of ~11.2 kb and were used for the main downstream analyses (Supplemental table 1). The longest read that passed the QC filtering in this dataset version was ~148.4 kb.

**Figure 1.**
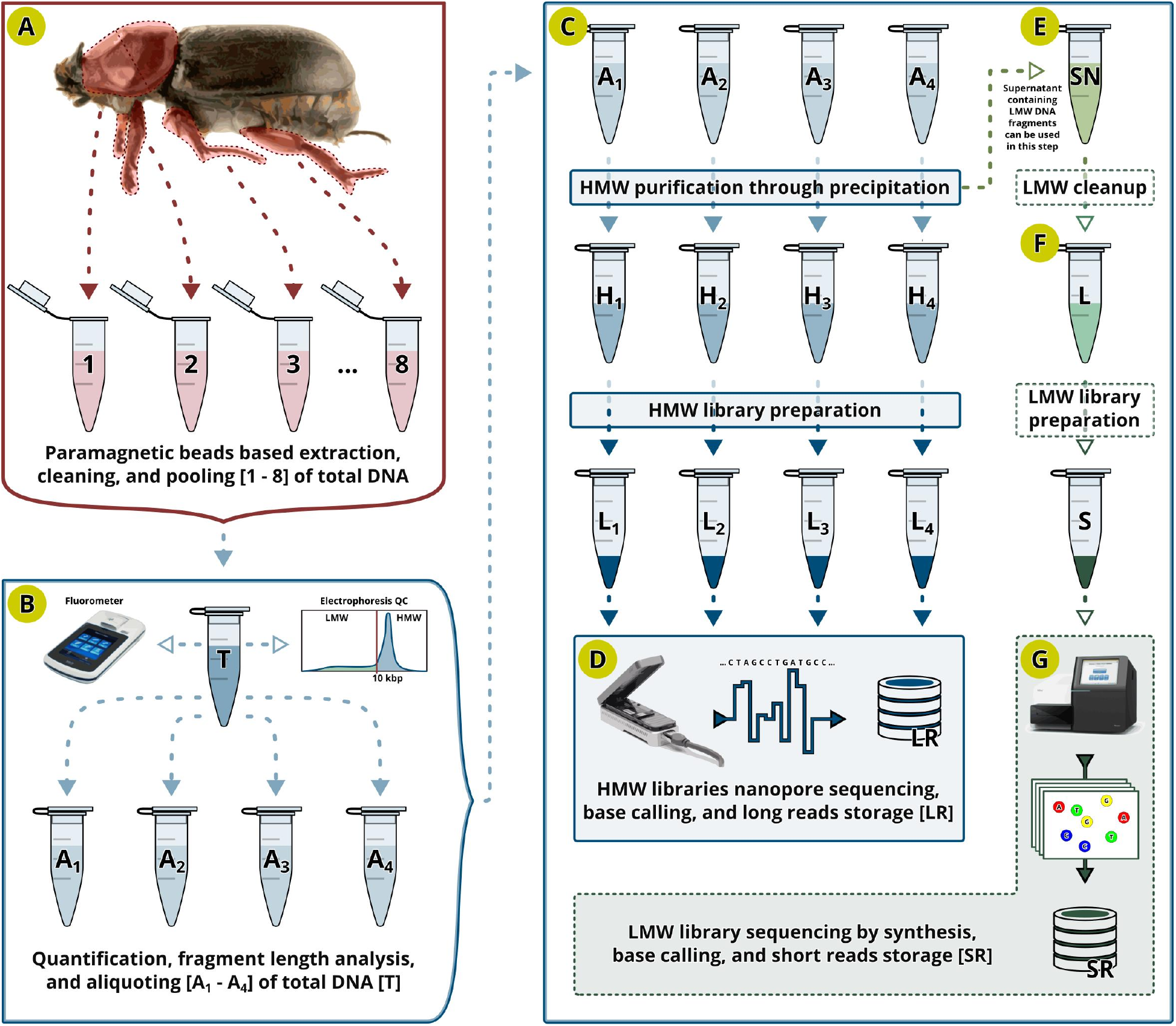
(A) Total DNA is extracted from thorax and legs using a bead-based protocol in 8 tubes, and all extracts are pooled (B) and assessed for DNA quality and quantity (electrophoresis QC, fluorometer). Two micrograms of DNA are aliquoted in each of four tubes [A1-A4] and (C) high molecular weight [HMW] DNA is precipitated [H1-H4] and used for HMW nanopore library preparations. (D) Each library [L1-L4] is sequenced on one Oxford Nanopore MinION flow cell. High accuracy base caller is used to transform the raw nanopore data into long reads [LR] and store them into fastq files for further analysis. (E) Supernatant [SN] containing low molecular weight DNA [LMW] can also be cleaned [L] and used for the preparation of LMW library [S] and high accuracy short read sequencing [SR].

### Genome assembly and quality assessment

Because we expected the long-read data (LR) to contain some percentage of mitochondrial, bacterial and other contaminant DNA reads, we first ran the long-read assembler Flye version 2.5 in metagenome mode that accommodates a highly non-uniform coverage of genomic fragments and is sensitive to under-represented sequences [27]. The initial draft assembly graph (S4-i-v1-g, Figure 2B) consisted of 512 nodes with N50 length of 7.9 Mb and total assembly size of 370.36 Mb. This initial draft assembly graph was then screened for the mitochondrial genome sequence, expecting a circular node 11 kb to 22 kb in size (based on a typical mitogenome size in insects [28]), and a disproportionately high depth of coverage (given that there are tens/hundreds of copies of the mitochondrial genome per nuclear genome copy in each cell). We identified one node with such characteristics: edge_110 (Figure 2D) was 21,039 bp in length and had a median coverage of 10,292X, showing the NCBI ‘blastn’ match with the mitochondrial genome assembly sequences (complete or partial) of beetles and other insects. Another circular node (edge_371) (Figure 2C) with a high depth of coverage (1,196X) was 126,204 bp in length, which we identified through the NCBI ‘blastn’ search as the Oryctes rhinoceros nudivirus (OrNV), a double-stranded DNA virus used as a biocontrol agent against *O. rhinoceros* [29]. Both nodes were removed from the draft assembly graph and analysed separately (Figure 2E-F), and their detailed characterization is described elsewhere [25], [26].

**Figure 2.**
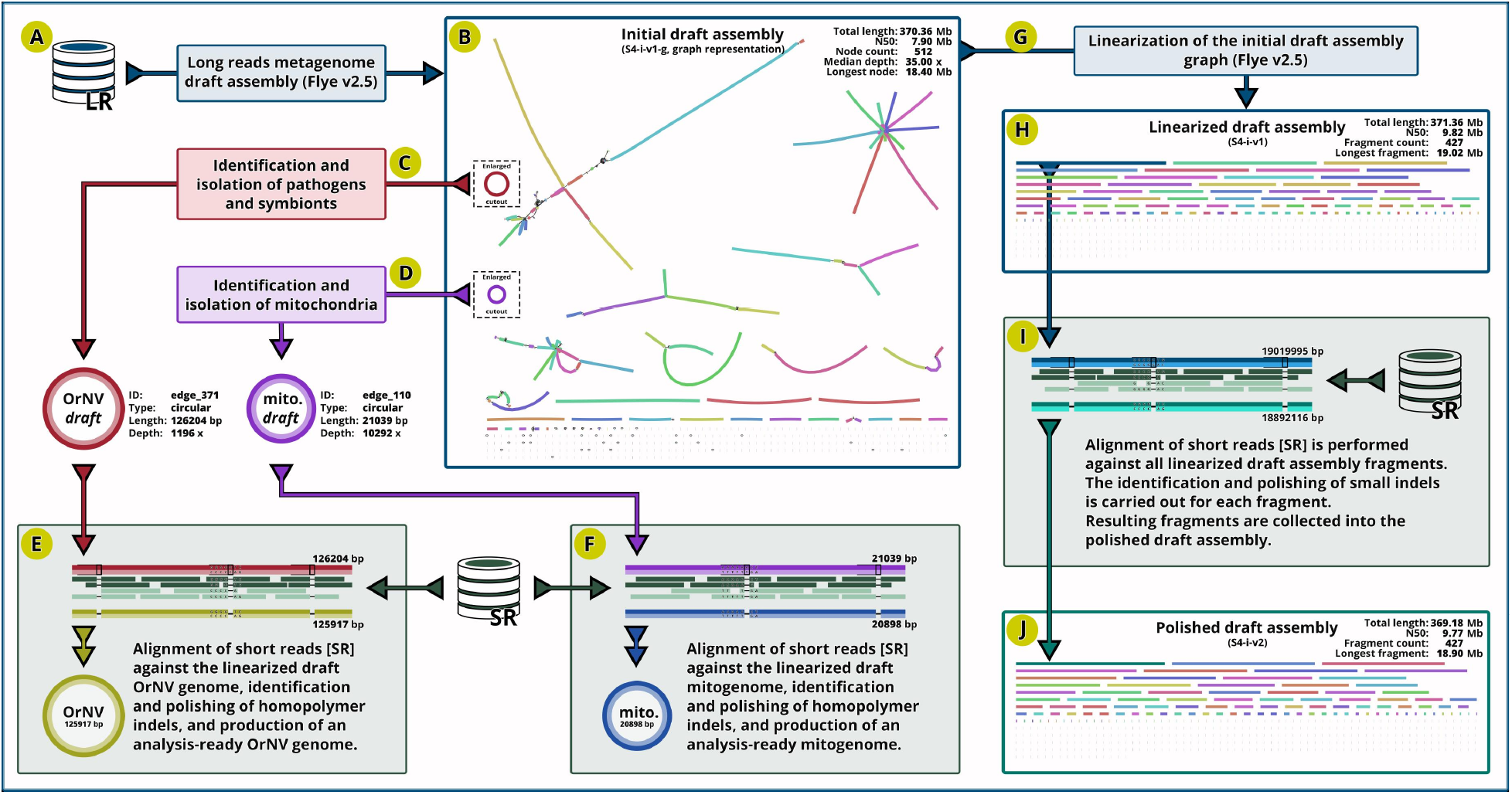
(A) Long-read data [LR] were used to generate the initial draft assembly (B), from which we identified and extracted the circular assembly for OrNV (C) and mitochondria (D). Short-read data [SR] were used to remove erroneous indels in homopolymers (E-F) to produce analysis-ready assemblies [25], [26]. The remainder of the draft assembly (G) was linearized (H) and short reads [SR] were used to remove erroneous indels (I) in each scaffold, producing an initial polished nuclear genome assembly for *Oryctes rhinoceros* (J).

Given the potential for ONT basecalling to introduce systemic indel errors in the homopolymer regions of the ONT-based assemblies [30], we used Pilon [31], BWA-MEM aligner [32] and more accurate Illumina Whole Genome Sequence data to remove small indels in the initial draft assembly (Figure 2H-I). We used the previously generated Illumina short reads from a whole-genome sequencing library that we prepared using the NebNext Ultra DNA II Kit (New England Biolabs, USA) with DNA extracted from another *O. rhinoceros* female that was collected from the same geographic location. The short-fragment Illumina library (Figure 1F-G) contained ~39.4 Gb of 150 bp paired end read data. We point out that Illumina sequencing library intended for the polishing of an ONT-based assembly would ideally be prepared from the same individual that was used to generate the long-read data. This would allow not only the correction of indels but also the correction of SNPs in the assembly consensus sequences. For the experiments with small-bodied insects that yield limited amounts of DNA, we recommend using the Low Molecular Weight (LMW) DNA found in the supernatant of the ONT library preparation mix (LMW depletion step, Figure 1E).

It is also worth noting that the indel error correction with the Illumina short reads has limitations in repetitive regions of the assembly, where short reads cannot be accurately aligned. For polishing, we used 92.4% of the Illumina reads that aligned to the initial genome assembly (S4-i-v1). Of the remaining reads, 6.1% aligned to the mitogenome and 0.2% to the OrNV genome, leaving 1.3% of the short-reads unaligned. The resulting polished initial genome assembly version S4-i-v2 (Figure 2J) consisted of 427 fragments (6 scaffolds and 421 contigs), with the fragment N50 length of 9.77 Mb, the longest fragment of 18.9 Mb, and a total assembly size of 369.18 Mb (34.9% GC content).

A quantitative assessment of the initial assembly’s accuracy and completeness was done through the benchmarking analysis of conserved genes, as implemented in BUSCO [33]. Using the BUSCO collection of 2,124 genes from the endopterygota database (endopterygota_odb10), we found that the initial polished assembly (S4-i-v2) contained 97.88% complete genes, with 97.18% occurring as single copies and only 0.94% missing. In comparison, BUSCO analysis of the unpolished assembly version (S4-i-v1) recovered only 65.1% genes as complete and 19.6% as missing, revealing the substantial impact of the uncorrected indel errors on gene prediction and detection (Supplemental table 2).

To further improve the assembly quality, we used the latest available version of the base-caller Guppy (v4.2.2) in high accuracy mode, and the latest available version of the long-read assembler Flye (v2.8.2) to generate multiple draft assemblies (Figure 3A-B) by increasing the minimum read overlap parameter for each assembly from 5 kb to 10 kb in increments of 500 bases. The Illumina short-reads were aligned against each draft assembly using BWA-MEM (Figure 3C), and the resulting alignments were further utilised to polish indels within each draft assembly (Figure 3D). This iterative process produced a collection of 11 polished draft assemblies (Figure 3E), and each was assessed for contiguity (assembly-stats “https://github.com/sanger-pathogens/assembly-stats”) and completeness (BUSCO) (Figure 3F) (Supplemental table 2). The best overall assembly (S4-7k-1v2) was produced with a minimal read overlap of 7 kb, and this parameter value was used to repeat the assembly, polishing and assessment two additional times (producing S4-7k-2v2 and S4-7k-3v2). The best of these three versions (S4-7k-2v2) was selected for further processing. We then removed the OrNV and mitochondrial sequences from the assembly (published previously [25], [26]), and this version (S4-7k-2v3) was further analysed with DIAMOND [34] and MEGAN [35], [36] in order to identify potential contaminant fragments. All assembly sequences that were not classified within Arthropoda in this pipeline were additionally checked against the NCBI’s online databases of nucleotide (nt/nr) and non-redundant protein sequences (nr) to identify the origin of a putative contaminant sequence (Figure 3G). Given that none of the analysed sequences had a significant BLAST hit to a taxon other than Coleoptera, we did not consider them as contaminants and did not remove them from the final genome assembly (S4-74-2v3, Figure 3H). This final assembly consisted of 1,013 fragments (6 scaffolds and 1,007 contigs), with the fragment length N50 of 10.70 Mb and the longest fragment (contig_6) of 32.67 Mb (Supplemental table 2).

**Figure 3.**
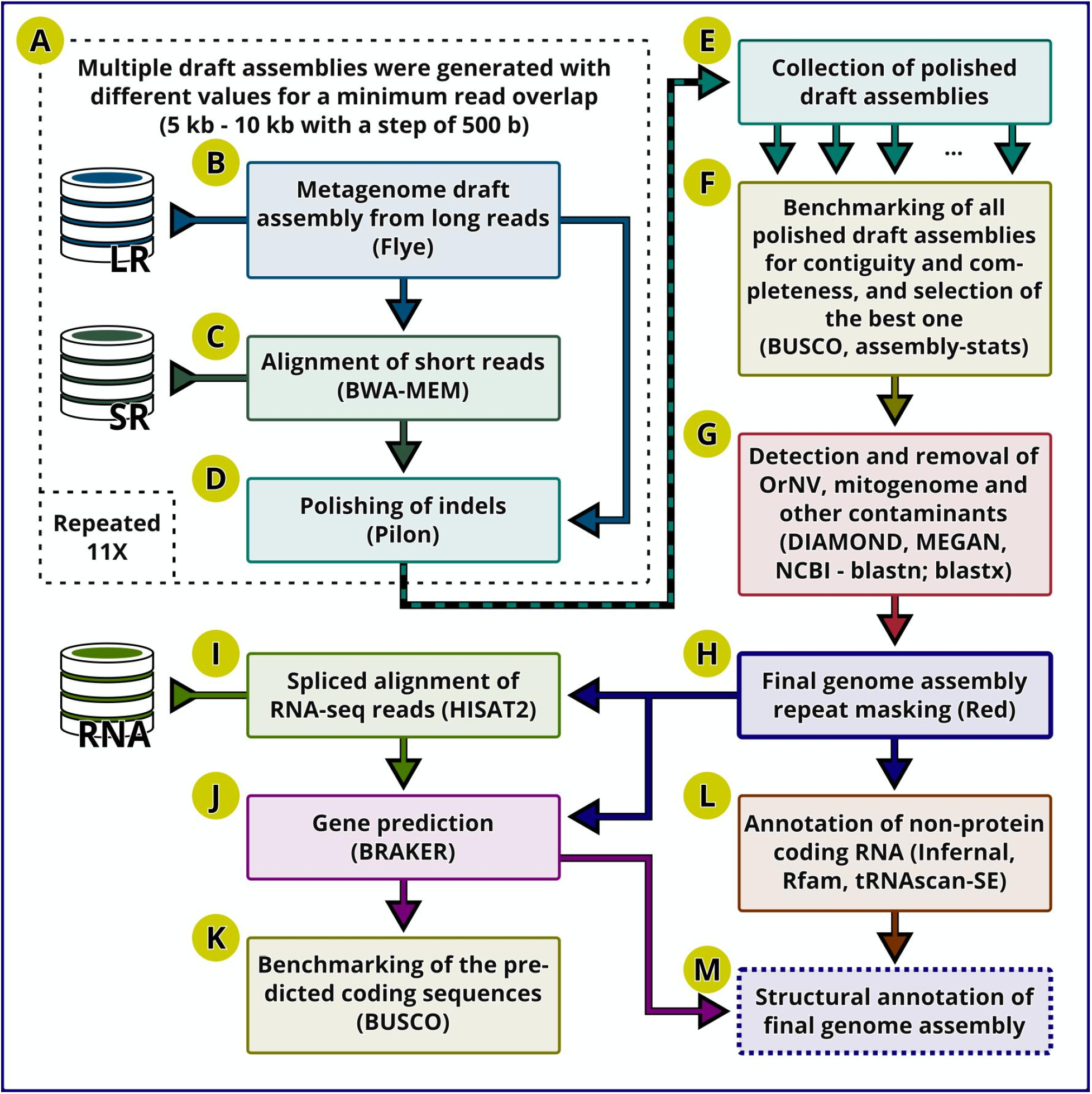
(A) Multiple polished draft assemblies were generated (B-D), collected (E) and benchmarked for completeness and contiguity (F) in order to determine the optimal read overlap for the long lead [LR] dataset. (G) Optimal draft assembly was screened for potential contaminants after the OrNV and mitogenome were removed. (H) The repeats were detected and soft masked in the final genome assembly. The splice-aware alignments (I) of the RNAseq datasets [RNA] were used for the prediction of protein coding genes (J), which was then assessed for completeness (K). Annotations of the non-protein coding RNAs (L) were added to form the final structural annotation of the nuclear genome assembly (M).

The size of our final *O. rhinoceros* nuclear genome assembly (S4-7k-2v3) was 377.36 Mb, which is very similar to the latest assembly for the congeneric beetle *O. borbonicus* (371.60 Mb in ungapped length, NCBI accession: GCA_902654985.1). The quality of our *O. rhinoceros* assembly, however, is superior to that of *O. borbonicus*, both in terms of contiguity (contig L50: *O. rhinoceros* vs. *O. borbonicus* = 12 vs. 571 (Supplemental table 3)) and completeness (BUSCOs: *O. rhinoceros* = 99.11% complete, 0.47% missing; *O. borbonicus* = 96.05% complete, 3.53% missing) (Supplemental table 4). Of note is that the original assembly for *O. borbonicus*, generated with the short-read Illumina technology, was first reported to be 518 Mb [37], but refinement with the 10X Genomics data led to a 28% reduction in size (removal of more than 140 Mb). The inflated size of the initial assembly was explained as a consequence of an incorrect haploidization of the assembly i.e., divergent haplotypes were assembled separately across many parts of the genome [38]. This exemplifies the difficulties of the assembly process based on the short-read sequencing of samples that have high genome-wide variability. Conversely, our *O. rhinoceros* assembly indicates that the correct haploidization is not problematic for long-read assemblers like Flye [27], particularly when the long-read data are generated from a single insect.

### Comparison with other available nuclear genome assemblies in Coleoptera

A recent ‘state of the field’ overview of insect genome assemblies [18] reports that this biological resource has been significantly underrepresented in Coleoptera (i.e. few genome assemblies are produced relative to the species richness), but that long-read sequencing is revolutionizing the creation of high-quality assemblies across insect groups [18]. We analysed 39 representative nuclear genome assemblies in the Coleoptera (out of 41 accessed from NCBI’s GenBank in October 2020) and found that one third were generated with data that included long-read sequences (nine assemblies with PacBio, four with ONT). For a total set of 39 analysed assemblies (Figure 4), the mean fragment N50 was 6.9 Mb (median: 298.9 kb, SD: 19.9 Mb) and the mean BUSCO completeness was 88.4% (median: 92.4%, SD: 14.27%). These quality metrics are above the average for a set of 601 assemblies from 20 insect orders (N50: 1.09 Mb, BUSCO completeness: 87.5%, [18]).

**Figure 4.**
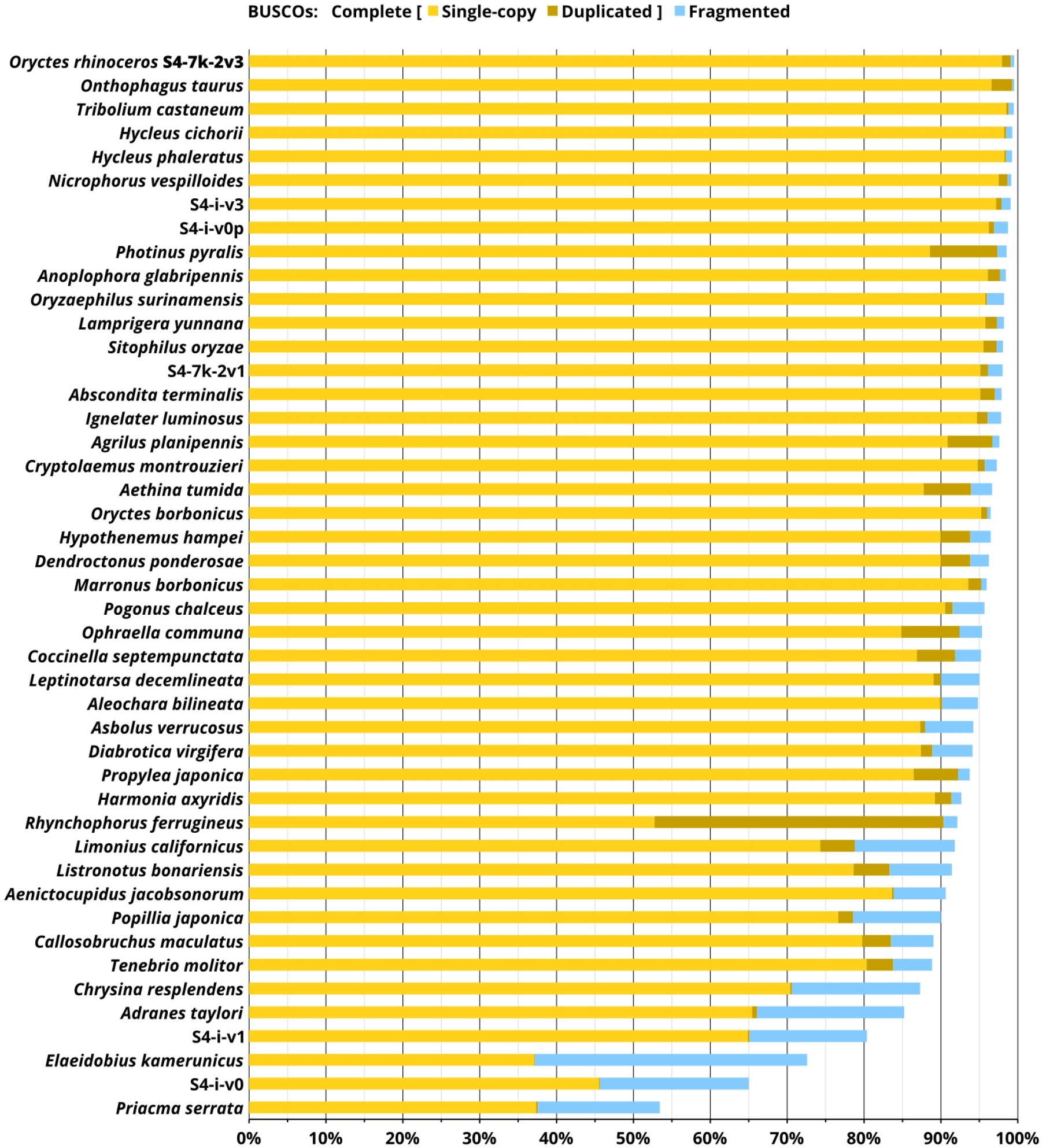
Comparison between multiple genome assemblies for *O. rhinoceros* and other available genome assemblies in Coleoptera, ranked by the number of detected BUSCOs. Assemblies of the highest quality have a very high percentage of complete single copy BUSCOs (yellow bar), and a small percentage of duplicated BUSCOs (brown bar) as well as fragmented BUSCOs (blue bar). *Oryctes rhinoceros* assemblies are: S4-7k-2v3 (final polished version), S4-i-v3 (initial polished version), S4-i-v0p (draft polished version from a single flow cell of long-read data), S4-7k-2v1 (final unpolished version), S4-i-v1 (initial unpolished version), S4-i-v0 (draft unpolished version from a single flow cell of long-read data).

Our *O. rhinoceros* assembly had the highest assembly accuracy and completeness among 39 published Coleopteran genomes, having only 0.47% missing BUSCOs (10 out of 2,124 core genes) and 0.42% fragmented BUSCOs (9 out of 2,124 core genes) (Figure 4). A genome assembly from another member of the family Scarabaeidae, *Onthophagus taurus*, had the same number of missing BUSCOs but twice as many duplicated genes (2.73%), and a substantially lower assembly contiguity, with scaffold (fragment) L50 of 160 versus 12 in *O. rhinoceros* (Supplemental Table 4). The improvements in the later versions of both the ONT basecaller Guppy and the long-read assembler Flye were reflected in a substantially better draft assembly of the genome prior to any indel polishing (see Figure 4: S4-7k-2v1 versus the equivalent nonpolished assembly S4-i-v1 that was produced with the older software versions).

### Structural annotation and quality assessment of the assembly

To delineate protein-coding genes, we used the BRAKER pipeline (Figure 3J) which enables an automated training of the gene prediction tools (GeneMark-EX and AUGUSTUS) with the extrinsic evidence from the RNA-Seq experiments [39]–[45]. We used the publicly-available RNA-seq data that cover different life stages of *O. rhinoceros*, from early instar larva, late instar larva, pupa, and the adult stage (NCBI accession: PRJNA486419; [46]), which is expected to maximize the probability of capturing the sequences of the entire set of expressed genes in this organism. To check data quality from these RNA-seq samples, we first aligned the reads against our genome assembly with the splice-aware aligner HISAT2 [47], and used these alignments to produce a genome-guided transcriptome with Trinity [48]. The assembled transcriptome had a very high BUSCO completeness (97.5%), indicating that the source RNA-seq dataset provides an excellent training set for gene prediction. Along with these aligned RNA-seq reads, the BRAKER pipeline was supplied with the final genome assembly (S4-7k-2v3) that had the repetitive regions (transposons and simple repeats) soft-masked on 32.73% of the assembly sequences (using the repeat detector Red [49]). The gene prediction algorithm produced a set of 16,375 protein-coding genes with a total of 20,072 transcripts. Our results match the available data for other members of Coleoptera; for example, 16,538 genes were reported for the bull-headed dung beetle *Onthophagus taurus* (Scarabaeidae) [50], [51], and the latest reference annotation for the red flour beetle *Tribolium castaneum* (Tenebrionidae) reports 16,593 genes with a total of 18,536 transcripts [52].

The benchmarking analysis (Figure 3K) indicated that our structural annotation of protein-coding genes in *O. rhinoceros* assembly is of high quality, showing BUSCO completeness of 92.09%. Somewhat lower completeness values obtained for the annotated gene set when compared to the assembly (92.09% vs. 99.11%) could indicate that the annotation pipeline, which uses multiple sources of evidence, has generated slightly inferior gene models for a set of single-copy orthologs than the single-predictor approach that BUSCO takes when working directly on the assembly sequences [53]. Such differences have been reported, for example, in the BUSCO assessment of 15 Anopheles mosquito genomes and their annotated gene sets [53].

The BUSCO metrics are based on the analysis of universal single-copy orthologous genes, but we wanted to check if paralogous genes are also correctly predicted in our annotation. Sigma Glutathione-S-Transferase genes (Sigma GSTs) belong to an ancient gene family and one of six classes of cytosolic GSTs in insects, that were reported to have undergone Oryctes-specific expansion [37]. Meyer and colleagues found 12 Sigma GST paralogs in the their *O. borbonicus* assembly, while the genomes of four other insects they analysed, including two beetles (*Tribolium castaneum* and *Dendroctonus ponderosae*), did not have more than seven paralogues in this GST class [37]. Based on this pattern, they hypothesized that the expansion of Sigma GST genes occurred specifically in the beetle lineage containing Oryctes species. However, our data do not support this hypothesis. Namely, we identified seven Sigma GST genes in two clusters in our *O. rhinoceros* annotation (Supplemental file 1), and this matches the structure in the red flour beetle, *T. castaneum* [54]. Considering that the initial *O. borbonicus* assembly contained divergent haplotypes that were not correctly haploidized [38], it was difficult (if not impossible) for Meyer and colleagues to avoid erroneous inference of gene duplications. Our *O. rhinoceros* assembly, on the other hand, enabled the annotation of the Sigma GST gene family that is consistent with other Coleoptera; moreover, the proteins they code show the highest amino-acid sequence similarity to the proteins of another scarab beetle (*O. taurus*, Figure 5). As expected for a GST family [54], Sigma GST genes in *O. rhinoceros* are located on one chromosomal section (contig_16) in two clusters, and like in *T. castaneum*, they have 3-5 introns and code 200-300 amino acids, except for one gene (K3A94_g10078) that has 12 introns and codes 434 amino-acids.

**Figure 5.**
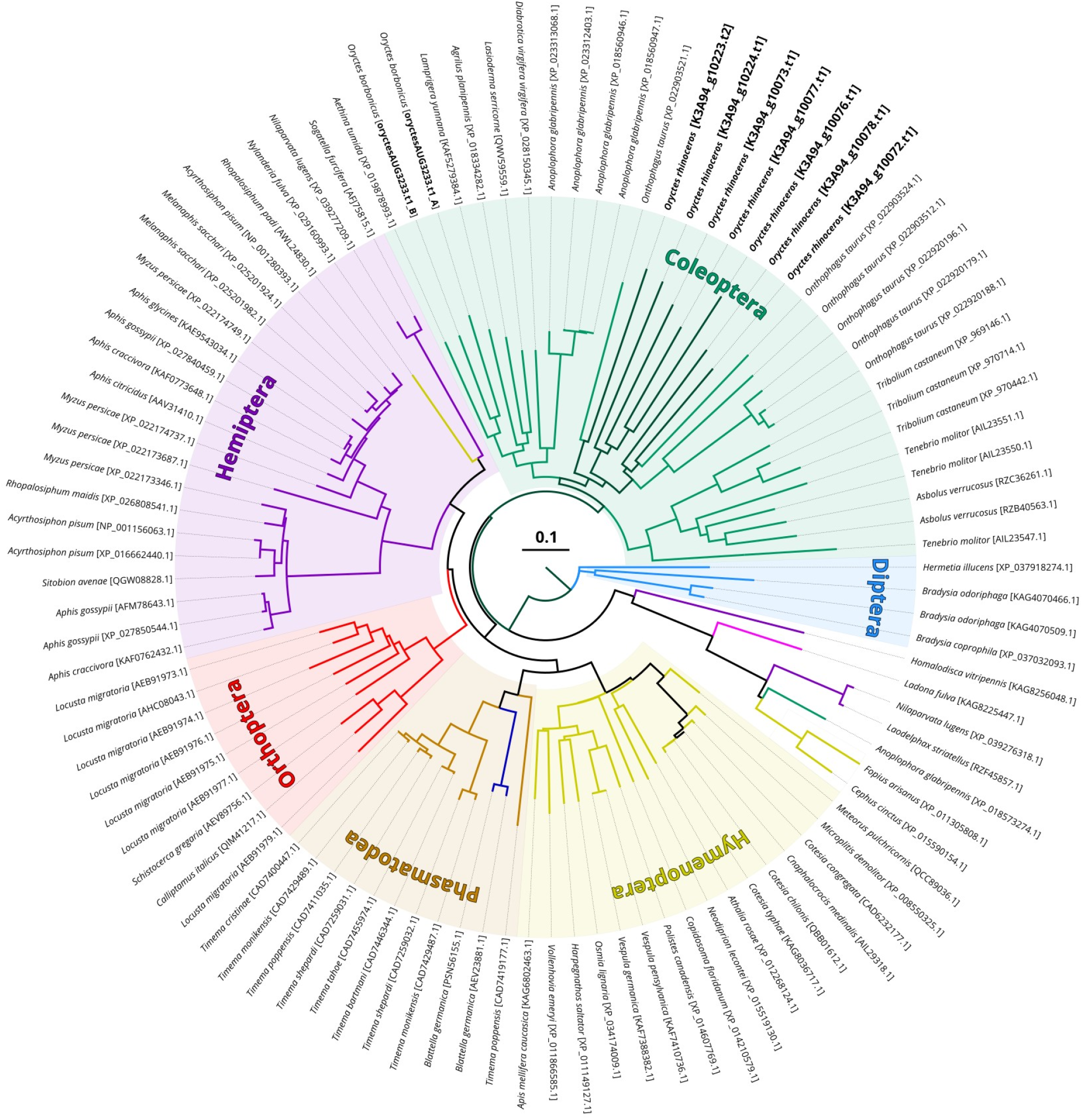
Neighbour joining (NJ) tree for protein sequences of putative Sigma GST genes identified in *O. rhinoceros* (dark green) and best BLASTp matches from the NCBI database. Distance between sequences (modelled as Grishin distance) used for tree generation predicts expected fraction of base substitutions per site given the fraction of mismatched bases in the aligned region. All branches (sequence groups) are based on >0.85 maximum sequence difference. Seven putative GST paralogs from *O. rhinoceros* show the greatest sequence similarity to another scarab beetle, *O. taurus*, and then other Coleoptera. With few exceptions, Sigma GST sequences from the members of the same order are grouped together (Coleoptera - green, Hemiptera - purple, Orthoptera - red, Phasmatodea - brown, Hymenoptera - yellow, Diptera - blue).

We also delineated the non-protein-coding RNAs, using tRNAscan-SE [55] and Infernal [56] with the Rfam database [57], [58] (Figure 3L). The annotation produced predictions for 18 tRNA-like pseudogenes, one selenocysteine tRNA gene, and 13 unknown isotypes. The number of tRNA genes predicted in *O. rhinoceros* (392) is highly congruent with another scarab beetle, *Onthophagus taurus*, that has 395 predicted tRNA genes [50], [51]. Our annotation with all predicted protein-coding genes, as well as non-protein-coding genes (including histones, rRNA, miRNA, …) and other features is provided as a gff3 file (Figure 3M) (Supplemental file 1).

### Application of genomic resources for management of *O. rhinoceros*

The assembled and annotated genome of *O. rhinoceros* provides an excellent opportunity to get genome-wide insight into the interaction between this insect pest and its control agent, Oryctes rhinoceros nudivirus (OrNV). For example, differences in the pattern on genome-wide expression can be traced between insects that have been experimentally infected with OrNV and the control group (non-infected) via transcriptome analysis. This approach for identifying putative infection-responsive genes has been used to study the interaction between one of the most important crop pests, the diamond-back moth *Plutella xylostella*, and the fungal insect pathogens, *Beauveria bassiana* and *Metarhizium anisopliae*, that have been widely used as insecticides [59]. For this type of a study, having access to a high-quality genome annotation is very important, as it has been shown that quality of a genome annotation strongly influences the inference of gene expression [60]. Identifying key *O. rhinoceros* genes that respond to OrNV infection could narrow a search for the causal genomic changes underlying the suspected attenuation of OrNV pathogenicity against this beetle. Namely, the resurgence and spread of *O. rhinoceros* over the last decade is hypothesized to be driven by the emergence of the virus-tolerant beetle populations and/or less virulent OrNV strains. The molecular basis for this suspected change in the beetle-OrNV interaction could reside in the regulation of small interfering RNAs (siRNAs) that are a known part of the insect immune response to viral infections [8], [61]. Our annotation contains the predictions for various non-protein-coding RNAs, laying a good foundation for further in-depth characterization of these regulatory genomic elements in *O. rhinoceros*.

RNA interference (RNAi) is a promising new approach for insect pest control, particularly for beetles that exhibit a robust environmental RNAi response [62], [63]. RNAi is a highly-specific gene-silencing mechanism that, when targeting essential insect genes, causes rapid mortality and could be developed into a control tool that is integrated with other pest management tactics. Our genome annotation facilitates the discovery of highly efficacious RNAi targets in *O. rhinoceros* through the mining of genes that are orthologs of the experimentally-validated targets in other beetles, such as *T. castaneum* and *D. v. virgifera* [64].

## CONCLUSIONS

We provide a highly contiguous and accurate nuclear genome assembly and structural annotation for an important invasive pest of palm trees, the scarab beetle *O. rhinoceros*. The assembly is based on the ONT sequencing of a single wild female, further demonstrating the utility of long-reads (and ONT sequencing in particular) in generating high-quality *de novo* genome assemblies from field specimens. Along with our structural annotation, this genomic resource opens up avenues for further biological discoveries aiming to improve the management of this pest, from the functional studies of interactions with the existing biocontrol agents, to the development of new control solutions *via* RNAi tools.

## MATERIALS AND METHODS

### Field collection and DNA isolation

*Oryctes rhinoceros* adults were collected from a pheromone trap (Oryctalure, P046-Lure, ChemTica Internacional, S. A., Heredia Costa Rica) on Guadalcanal, Solomon Islands in January 2019 and preserved in 95% ethanol. High-molecular weight (HMW) DNA was extracted from a single female using a customized paramagnetic (SPRI) bead-based protocol. Specifically, we dissected pieces of tissue from four legs and the thorax, avoiding the abdomen to minimize the proportion of gut microbiota in the total DNA extract (Figure 1A). We incubated approximately 50 mm^3^ of tissue in each of the eight 1.7 mL eppendorf tubes with 360 μL ATL buffer, 40 μL of proteinase K (Qiagen Blood and Tissue DNA extraction kit) for 3h at room temperature, while rotating endover-end at 1 rpm. Four hundred microliters of AL buffer were added to each reaction and incubated for 10 min at room temperature, followed by the addition of 8 μL of RNase A and incubation for 5 minutes at room temperature. To remove the tissue debris, each tube was spun down for 1 min at 16,000 rcf and 600 μL of homogenate was transferred to a fresh tube. Six hundred microliters of the SPRI bead solution were added to each homogenate and incubated for 30 min while rotating at end-over-end at 1 rpm. After two washes with 75% ethanol, DNA in each tube was eluted in 50 μL of TE buffer. All eight elutions were combined and DNA quality was assessed on the 4200 Tapestation system (Agilent) and with the Qubit broad-range DNA kit (Figure 1B). Finally, we used the Circulomics Short Read Eliminator XS kit to enrich the DNA elution with fragments longer than 10 kb (i.e. High Molecular Weight, HMW, DNA, Figure 1C).

### ONT library preparation and sequencing

One microgram of the size-selected HMW DNA was used as the starting material for the preparation of each ONT library, following the manufacturer’s guidelines for the Ligation Sequencing Kit SQK-LSK109 (Oxford Nanopore Technologies, Cambridge UK). Four libraries were sequenced on four R9.4.1 flow cells using the MinION sequencing device and the ONT MinKNOW Software (Oxford Nanopore Technologies, Cambridge UK) (Figure 1C).

### Genome assembly

High accuracy base-calling from the raw ONT data was computed with Guppy v3.2.4 (for the initial assembly) and Guppy v4.2.2 (for the final assembly). The initial genome assembly (S4-i-v1) was produced with Flye version 2.5 [27] using the following input parameters: the approximate genome size ( --genome-size) of 430 Mb, based on the size of an initial genome assembly in a related species *O. borbonicus* [37] two iteration of polishing ( --iterations 2), aimed at correcting a small number of extra errors based on the improvements on how reads align to the corrected assembly; a minimum overlap between two reads ( --min-overlap 5000) of 5000 bp; and a metagenome mode ( --meta) to allow for the recovery of mitochondrial, symbiont, pathogen and other “contaminant” genomes, given that this mode is sensitive to highly variable coverage and under-represented sequences [27]. Flye version 2.8.2 was used during the iterative process for the final genome assembly (S4-7k-2v), with the parameter ‘--min-overlap’ ranging from 5000 bp to 10000 bp in 500 bp increments while keeping other parameters (--genome-size, --iterations, --meta) unchanged.

### Identification of pathogens, symbionts, contaminants

Screening of the circular nodes with a disproportionately high coverage in the initial genome assembly graph identified the OrNV and mitogenome, and they were removed from further analyses. A linearized set of the remaining putative genome assembly sequences (contigs and scaffolds) were locally compared against the NCBI non-redundant protein (nr) database using DIAMOND [34] version 0.9.24 in ‘blastx’ mode. The NCBI database was downloaded from ftp.ncbi.nih.gov/blast/db/FASTA/. The results obtained with DIAMOND were analysed with the metagenome analyser tool MEGAN [36]. Any sequence not classified within Arthropoda was also checked against the NCBI’s online database of nucleotide (nt/nr) and non-redundant protein sequences (nr) to identify the origin of a suspected contaminant sequence.

### Polishing of the genome assembly with Illumina reads

Indel errors in the homopolymer regions represent inherent basecalling errors of the ONT platform [30]. To remove putative indel errors in the draft assembly, we used the genome polishing program Pilon version 1.23 [31] that was supplied with the spliced-aware alignments of the Illumina reads from one whole-genome sequencing library. DNA for this Illumina library originates from a female beetle collected in the same location as the female used for the ONT sequencing. Because Illumina and ONT data did not come from the same individual, we only performed indels polishing. The Illumina sequences were produced on a HiSeq X10 platform by Novogene (Beijing, China) using the 150 bp paired-end chemistry, and were processed in Trimmomatic [65] to remove Illumina adapters, and trim and filter each read based on the minimum phred score of 20.

### Evaluation of genome assemblies

The completeness of the initial genome assembly (S4-i-v3) was evaluated using: (a) alignment of DNA-seq data, (b) alignment of RNA-seq data, and (c) the recovery of the benchmarked universal single copy orthologs (BUSCOs) [33]. We used the BWA-MEM aligner with default settings and recorded the percentage of mapped Illumina reads from the whole-genome sequencing dataset (Illumina DNA library described above) and four independently-generated RNA-seq datasets from the beetle’s four life stages [46](NCBI SRA Accession: PRJNA486419) that were combined prior to alignment with the beetle genome assembly. The number of recovered universal singlecopy orthologs (SCOs) was obtained using the “genome autolineage” mode in BUSCO version 4.0.6, that first searched the databases ‘eucariota_odb10’ (7 species, 255 SCOs), and ‘endopterigota_odb10’ (56 species, 2,124 SCOs). To perform the comparative benchmarking, the same BUSCO analysis was done for 39 representative assemblies in the Coleoptera out of 41 that were available in the NCBI’s GenBank in October 2020 (Supplemental table 4). Two Coleoptera genomes (for *Protaetia brevitarsis* GCA_004143645.1, and *Alaus oculatus* GCA_009852465.1) were excluded due to a persistent BUSCO analysis failure with their assembly files.

### Structural annotation

To perform the structural annotation of the final genome assembly, we used the independently-generated RNA-seq datasets from the beetle’s four different life stages (NCBI SRA Accession: PRJNA486419)[46]. The RNA-seq reads were pruned of the Illumina adapters and aligned against our genome assembly with the splice-aware aligner HISAT2 (Figure 3I). The quality and completeness of these RNA-seq data were assessed through the transcriptome assembly in Trinity version 2.10.0 [48], [66], using the default settings in two modes: *de novo* and genome-guided assembly. To avoid incorporating the extraneous RNA sequences into the *de novo* transcriptome assembly, we used only those reads that were mapped with HISAT2 [47] to our S4-i-v3 genome assembly. The completeness of each transcriptome assembly was evaluated with BUSCO, using the ‘auto-lineage’ mode. The final genome assembly (S4-7k-2v3) and the splice-aware alignments (from HISAT2) were used for the genome-guided transcriptome assembly using the BRAKER pipeline version 2.1.4 (https://github.com/Gaius-Augustus/BRAKER/releases/tag/v2.1.4). Annotation of the non-coding RNA genes was done with tRNAscan-SE version 2.0.6 [55], [67] and Infernal version 1.1.3 [56] against the Rfam database v14.2 [57], [58] that was available on Sep 7 2020 (ftp://ftp.ebi.ac.uk/pub/databases/Rfam/14.2/).

### Analysis of the Sigma GST gene group

Genes from the Sigma Glutathione-S-Transferase group were identified through the BLASTp analysis [68], using the protein sequences translated from our *O. rhinoceros* genome annotation. First, we found eight predicted protein sequences in our annotation that had a significant BLASTp match (E-value << e-5) with two reported GST protein sequences from *O. borbonicus* [37]. Two of the eight identified *O. rhinoceros* sequences were transcript variants from one gene, so we chose one of the variants for further analyses. We then ran BLASTp analysis (with default parameters) with one putative Sigma GST protein sequence from *O. rhinoceros* (K3A94_g10072.t1) to identify highly similar sequences within the NCBI’s database. The list of 100 best matches (including six remaining *O. rhinoceros* sequences, and two *O. borbonicus* sequences) with NCBI’s accession numbers is found in the Supplemental Table 5. The unrooted tree was generated by the NCBI’s algorithm using the neighbour joining method [69], maximum sequence difference >0.85 and the modelled protein distance by Grishin [70]. The final newick tree file is available in the Supplementlary Data (Supplemental file 1). The tree visualization was done in FigTree [71].

## Availability of supporting data

The *Oryctes rhinoceros* genome assembly S4-7k-2v3 and raw reads used in this study are available for download via NCBI [Bioproject: PRJNA752921]. Supporting material including functional annotation and interim genome assemblies are available for download from [DOI:xxxxx].

## Authors’ contribution

**Igor Filipović:** Conceptualization, Investigation, Formal Analysis, Software, Methodology, Data Curation, Visualization, Writing - Original Draft Preparation, Writing - Reviewing & Editing.

**Gordana Rašić:** Methodology, Resources, Supervision, Writing - Reviewing & Editing.

**James Hereward:** Data Curation, Writing - Reviewing & Editing.

**Maria Gharuka:** Resources.

**Gregor J. Devine:** Resources, Supervision, Writing - Reviewing and Editing.

**Michael J. Furlong:** Resources, Funding acquisition, Conceptualization, Project administration, Supervision, Writing - Reviewing and Editing.

**Kayvan Etebari:** Resources, Funding acquisition, Conceptualization, Supervision, Writing - Reviewing and Editing.

## Acknowledgments

This project was supported by the Australian Centre for International Agricultural Research funding (HORT/2016/185), the University of Queensland (UQECR2057321) and by core funds from the Mosquito Control Laboratory at QIMR Berghofer MRI.

