## Supplementary figures and images for "A high-quality *de novo* genome assembly based on nanopore sequencing of a wild-caught coconut rhinoceros beetle (*Oryctes rhinoceros*)"

### Supplemental Figure 1

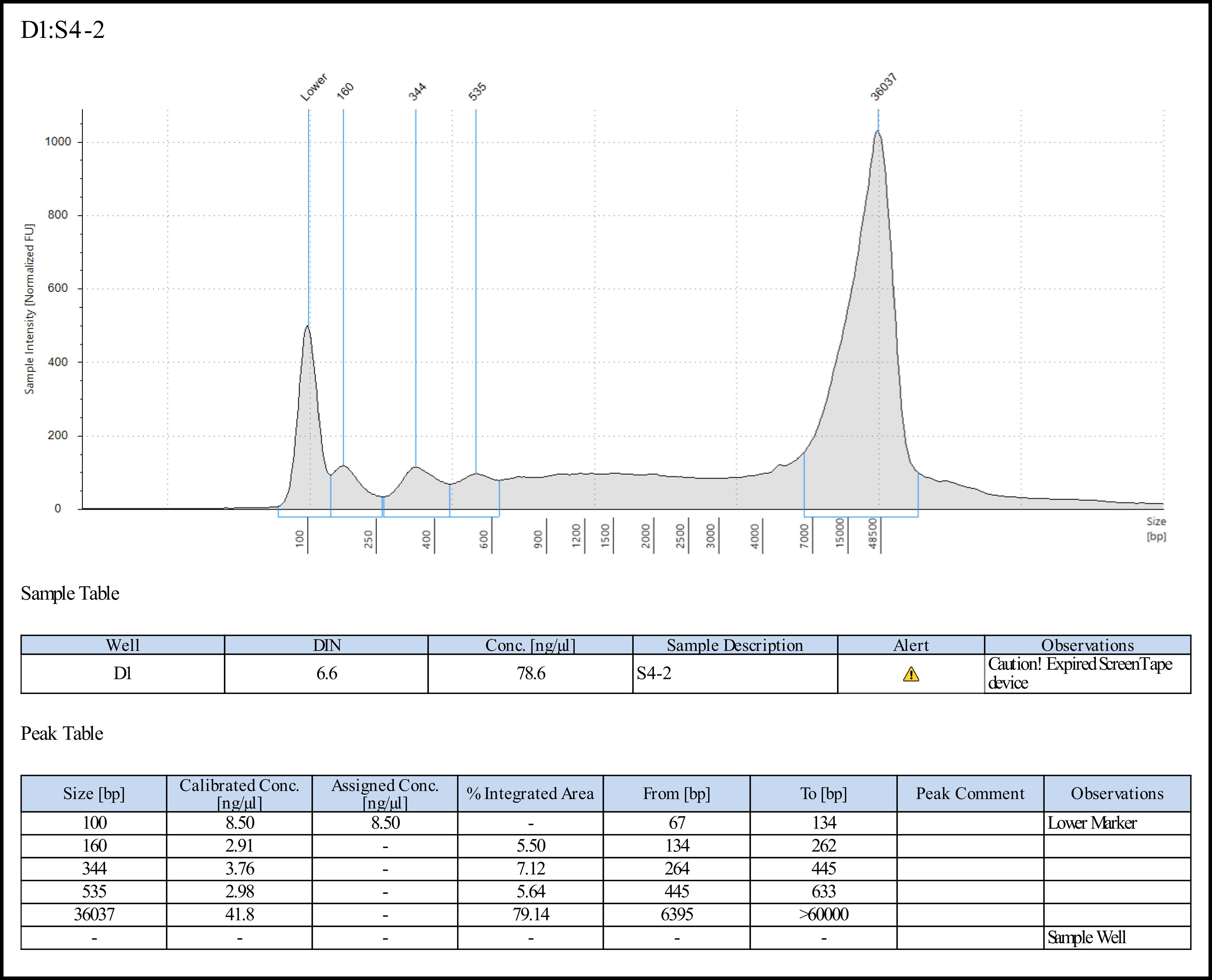
